# Glycogen-fuelled metabolism supports rapid Mucosal Associated Invariant T cell responses

**DOI:** 10.1101/2022.12.20.521164

**Authors:** Féaron C. Cassidy, Nidhi Kedia-Mehta, Ronan Bergin, Andrea Woodcock, Ardena Berisha, Ben M. Bradley, Eva Booth, Odhran K. Ryan, Linda V. Sinclair, Donal O’Shea, Andrew E. Hogan

## Abstract

Mucosal Associated Invariant T (MAIT) cells are a subset of unconventional T cells, which recognise a limited repertoire of ligands presented by the MHC class I-like molecule MR1. In addition to their key role in host protection against bacterial and viral pathogens, MAIT cells are emerging as potent anti-cancer effectors. With their abundance in human, unrestricted properties and rapid effector functions, MAIT cells are emerging as attractive candidates for cancer-immunotherapy. In the current study, we demonstrate that MAIT cells are potent anti-tumour cells, rapidly degranulating and inducing target cell death. Previous work from our group and others has highlighted glucose metabolism as a critical process for MAIT cell cytokine responses at 18 hours. However, the metabolic processes supporting rapid MAIT cell anti-tumour responses are currently unknown. Here, we show that glucose metabolism is dispensable for both MAIT cell cytotoxicity and early (<3 hours) cytokine production, as is oxidative phosphorylation. We show for the first time that MAIT cells have the machinery required to make and metabolize glycogen, and demonstrate that MAIT cell cytotoxicity and rapid cytokine responses are dependent on glycogen metabolism. In summary, we show for the first time that glycogen-fuelled metabolism supports rapid MAIT cell effector functions (cytotoxicity and cytokine production) which may have implications in their use as an immunotherapeutic agent.

## Introduction

Mucosal Associated Invariant T (MAIT) cells are a population of unconventional T cells which are important in the immune defence against bacterial and viral infections^1, 2, 3, 4, 5, 6^. MAIT cells are restricted by the MHC-like molecule MR1^5^, and recognise a limited set of bacterially derived antigens^7^. MAIT cells are primed to respond, and display an inherent “innateness” with higher levels of effector molecule mRNA at the steady than conventional T cells^8^. MAIT cells can be activated either via TCR triggering or innate cytokine stimulation, after which they are capable of producing a range of cytokines and lytic molecules including IFNγ and granzyme B^9, 10^. These rapid effector responses allow MAIT cells to initiate and amplify the immune response, as well as directly targeting infected or transformed cells^11, 12, 13, 14^. Robust anti-cancer responses and the ability to activate other anti-cancer cells^14^ paired with their absence of MHC restriction, has highlighted MAIT cells as an attractive candidate for immunotherapy^14, 15, 16^.

Several studies have identified tumour-infiltrating MAIT cells in primary and metastatic lesions^13, 17, 18, 19^, but often report diminished effector function including loss of key cytokines such as IFNγ^13, 17^. Therefore, it is important to fully understand the molecular pathways regulating MAIT cell effector responses. Our previous work has demonstrated that MAIT cells undergo metabolic reprogramming in order to provide the energy and biosynthetic intermediates needed to support their robust effector functions^20, 21^. We and others have demonstrated that human MAIT cells activated via their TCR for 18 hours favour exogenous glucose as their carbon source, and engage in glycolytic metabolism as their primary metabolic program^20, 22^. This is mediated by the activation of the critical metabolic regulators mTOR and MYC, which control the expression of nutrient transporters, and key enzymes involved in metabolism of glucose^20, 21^.

Currently, the metabolic requirements for rapid MAIT cell effector responses such as cytotoxicity are unknown and were the focus of the current study. We show that MAIT cells co-cultured with cancer cells rapidly (within 2 hours) degranulate and induce cell death. We demonstrate that these rapid responses are independent of glucose-fuelled glycolytic metabolism, and show for the first time that MAIT cells contain the machinery required to synthesize and metabolize glycogen. We demonstrate that MAIT cells have glycogen stores, and inhibition of glycogenolysis inhibits MAIT cell cytotoxicity and early cytokine responses, which may have implications for the therapeutic use of MAIT cells.

## RESULTS

### MAIT cell respond rapidly with target cell lysis and cytokine production

We first assessed the expression of MR1 expression on several human cancer cell lines, and identified the K562 myelogenous leukaemia cell line as MR1+,; furthermore we demonstrate that the addition of 5-ARU-MG increased the expression of MR1 on the surface of the cell (Figure 1A). Next, we demonstrate that MAIT cells respond to K562 cells by degranulating (CD107a expression) and this is significantly boosted with the addition of 5-ARU-MG (Figure 1B-C). To confirm that MR1 was required for MAIT cell degranulation we next blocked MR1 and observed reduced degranulation (Figure 1D-E). To build on these findings, and to confirm if MAIT cells can induce target cell death, we moved to a direct cytotoxicity assay, and demonstrate that MAIT cells can rapidly (within 2 hours) kill K562 cells, and in a dose-dependent manner (Figure 1F). In addition to cytotoxicity, we also show that MAIT cells can rapidly upregulate IFNγ expression in response to TCR stimulation (Figure 1G-H).

**Figure 1.**
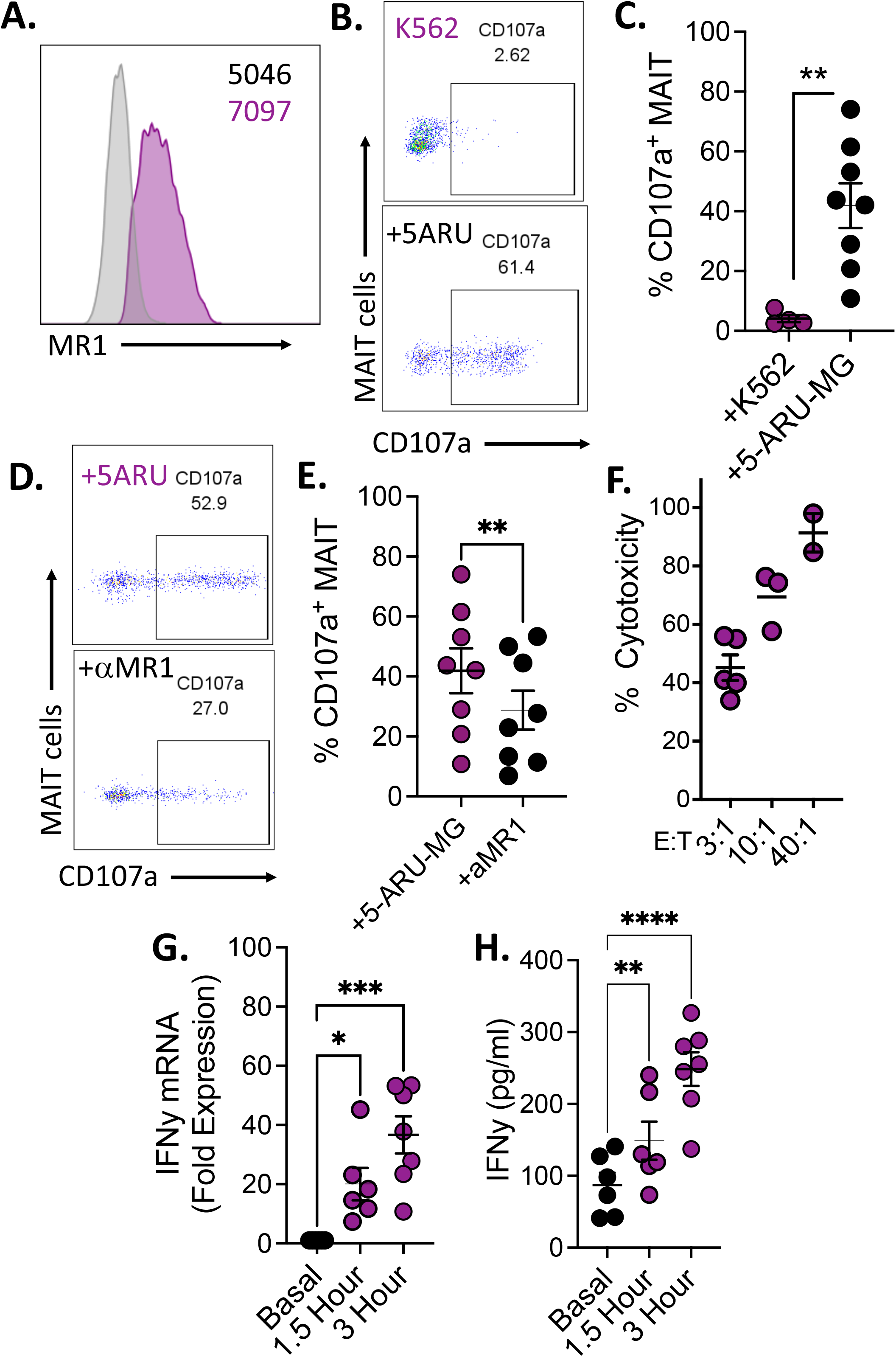
MAIT cell respond rapidly with target cell lysis and cytokine production. **(A.)** Flow cytometry histogram displaying MR1 expression on the surface of K562 cells in the absence (Grey) or presence (purple) of exogenous 5-ARU-MG (MFI annotated on plot). **(B-C)** Flow cytometry dot plots and scatter plot showing CD107a expression on *ex-vivo* MAIT cells cultured with K562 or K562 pulsed with 5-ARU-MG. **(D-E)** Flow cytometry dot plots and scatter plot showing CD107a expression on *ex-vivo* MAIT cells cultured with K562 pulsed with 5-ARU-MG in the absence or presence of MR1 blocking antibody. **(F)** Scatterplot showing dose-dependent (effector to target ratio) cytotoxicity of IL-2 expanded MAIT cells in their targeting of K562 cells pulsed with 5-ARU-MG. **(G-H)** Scatter plots of IFNγ mRNA and secreted protein levels from IL-2 expanded MAIT cells stimulated with TCR beads (antiCD3/CD28) for either 1.5 hours or 3 hours. ns = not significant, * = p > 0.05, ** = p < 0.01, *** = p < 0.001, **** = p < 0.0001 as measured by paired t test, Friedman test or mixed-effects analysis where appropriate.

### MAIT cell cytotoxicity is not dependent on glucose metabolism or oxidative phosphorylation

Our previously published data demonstrated that MAIT cells favour exogenous glucose as their carbon source, which is metabolised via glycolytic metabolism^21^. We and others have reported that glucose metabolism is critical for maximal MAIT cell IFNγ and granzyme production^20, 22^. To investigate if MAIT cell cytotoxicity was dependent on glucose metabolism, we utilized the glycolytic inhibitor 2-Deoxy-D-glucose (2DG) and show no effect on either MAIT cell degranulation or cytotoxicity (Figure 2A-C). Studies in other T cell subsets have demonstrated that expression of the glucose transporter (GLUT1) can take up to 6 hours after stimulation^23^. We therefore investigated expression of GLUT1 on TCR stimulated MAIT cells, and demonstrated that expression is not detectable at 3 hours post stimulation but is detectable at 6 hours (Figure 2D), further supporting the concept that rapid MAIT cell cytotoxicity is not supported by exogenous glucose metabolism. Additionally, we found that rapid upregulation of IFNγ was also not inhibited by addition of 2DG (Figure 2E). Another major metabolic pathway utilized by some T cell subsets is oxidative phosphorylation (OxPhos). To investigate if OxPhos supports MAIT cell cytotoxicity we utilized the specific inhibitor oligomycin and show no effect on either MAIT cell degranulation or lysis of target cells (Figure 2F-G).

**Figure 2.**
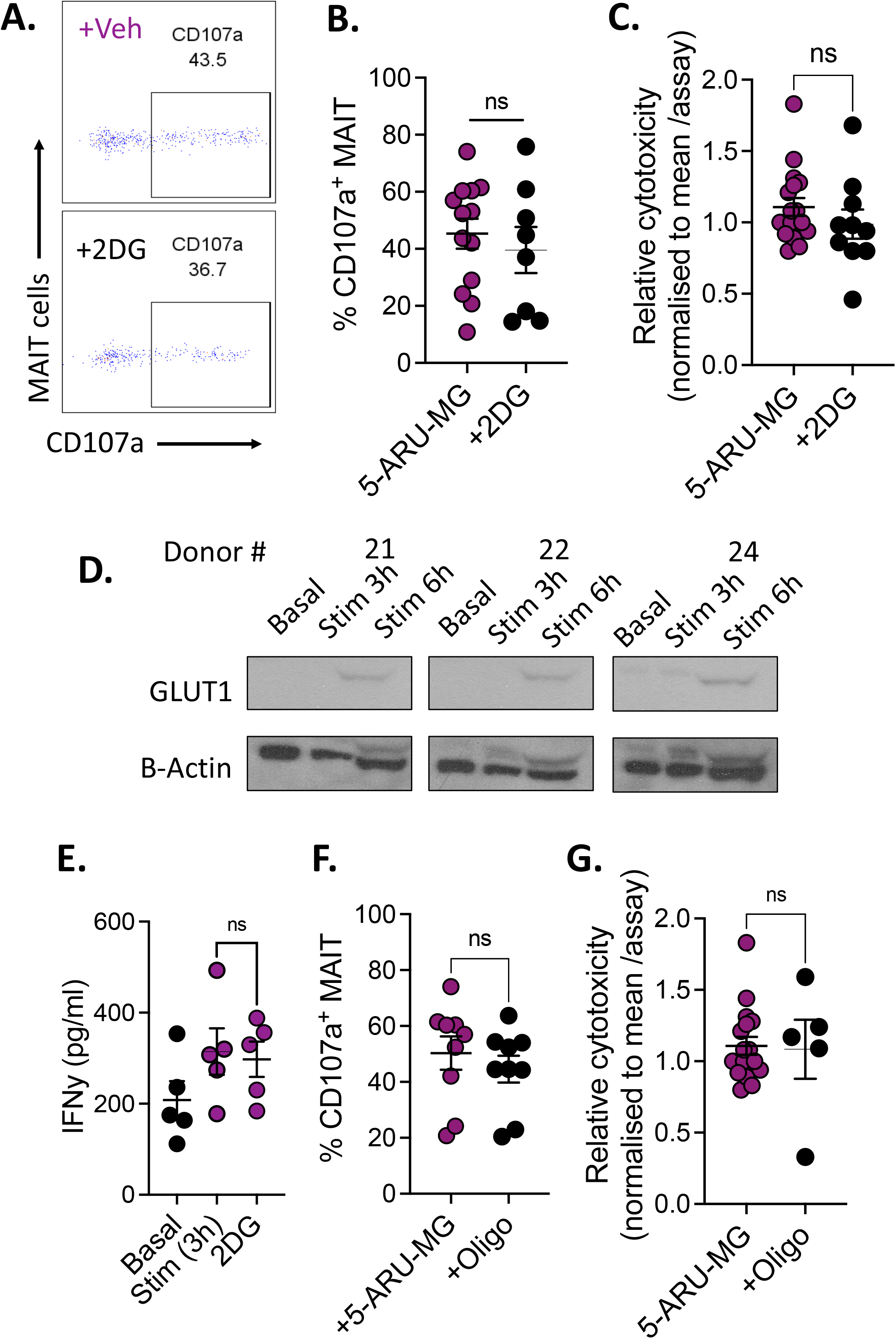
MAIT cell cytotoxicity is not dependent on glucose metabolism or oxidative phosphorylation. **(A-B)** Flow cytometry dot plot and scatter plot showing CD107a expression on *ex-vivo* MAIT cells in response to stimulation with K562 cells (pulsed with 5ARU-MG) with or without the addition of glycolytic inhibitor 2DG. **(C)** Scatterplot showing the impact of 2DG treatment on cytotoxic capacity of IL-2 expanded MAIT cells, displayed as fold change per sample. **(D)** Western Blot of GLUT1 protein expression by IL-2 expanded MAIT cells (3 individual donors) at rest (basal) and after 3hours or 6 hours of TCR bead stimulation. **(E)** Scatter plot showing the impact of 2DG treatment on early IFNγ secretion by IL-2 expanded MAIT cells stimulated with TCR beads. **(F)** Scatter plot showing CD107a expression on *ex-vivo* MAIT cells cultured with K562 (pulsed with 5ARU-MG) in the absence or presence of the oxidative phosphorylation inhibitor oligomycin. **(G)** Scatterplot the impact of oligomycin treatment on the cytotoxic capacity of IL-2 expanded MAIT cells, displayed as fold change per sample. ns = not significant, as measured by paired t test, Wilcoxon test or Mann-Whitney test as appropriate.

### MAIT cells contain the machinery to synthesize and metabolize glycogen

We next investigated other potential metabolic pathways which may support MAIT cell cytotoxicity by interrogating our publicly available MAIT cell proteomic dataset^21^ and identified that MAIT cells express the enzyme glycogen synthase (GYS-1) which is required to synthesize glycogen (Figure 3A). We next confirmed the expression via western blotting (Figure 3B-C). We investigated if MAIT cells expressed the enzymes required for the breakdown of glycogen and found the brain isoform of glycogen phosphorylase (PYGB) in our proteomics dataset and again verified via western blotting (Figure 3D-F). To investigate if PYGB was active in MAIT cells we utilized a glycogen phosphorylase activity assay and observed increased PYG activity in TCR stimulated MAIT cells (Figure 3G). Finally, we show that MAIT cells contain stored glycogen, and that upon stimulation glycogen is content is reduced (Figure 3H-I).

**Figure 3.**
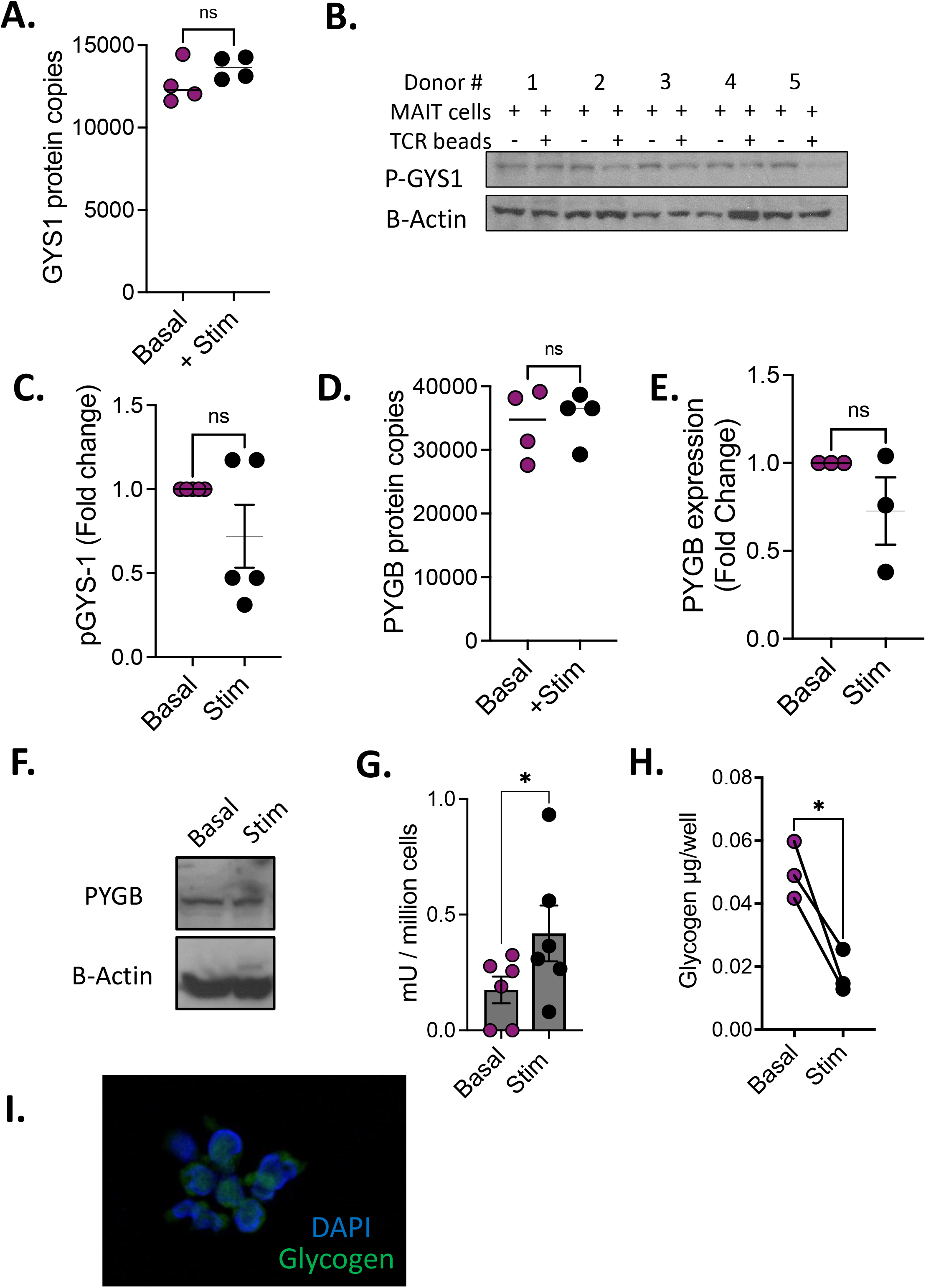
MAIT cells contain the machinery to synthesize and metabolize glycogen. **(A)** Scatter plot showing the expression of GYS1 in IL-2 expanded MAIT cells, as measure by proteomics. **(B-C)** Western blot and densitometry scatter plot showing the expression of phosphorylated GYS1 in MAIT cells. **(D)** Scatter plot showing the expression of PYGB in IL-2 expanded MAIT cells, as measure by proteomics. **(E-F)** Densitometry scatter plot and western blot showing the expression of PYGB (glycogen phosphorylase) in MAIT cells. **(G)** Scatter plot showing glycogen phosphorylase activity in TCR-stimulated MAIT cells, in the absence or presence of the glycogen phosphorylase inhibitor CP91149. **(H)** Scatter plot showing glycogen levels in IL-2 expanded MAIT cells at rest (basal) or stimulated with TCR beads for 3 hours. **(I)** Florescent microscopy image demonstrating the presence of glycogen in MAIT cells. ns = not significant, p > 0.05, * = p < 0.05 as measured by paired t test.

### Glycogen supports MAIT cells cytotoxicity and early cytokine responses

To investigate if glycogen supports MAIT cell anti-tumour responses we utilized the glycogen phosphorylase (PYG) inhibitor CP91149 and demonstrate the inhibition of MAIT cell degranulation (Figure 4A-B). To confirm, we utilized another glycogen phosphorylase inhibitor (GPI)^23^, and again observed reduced MAIT cell degranulation (Figure 4C). We next investigated if inhibiting the breakdown of glycogen limited target cell lysis by MAIT cells, and demonstrate a significant reduction in killing (Figure 4D). Another anti-tumour function of MAIT cells is their robust production of IFNγ. We investigated if early IFNγ cytokine production (<3 hours) is dependent on the metabolism of glycogen and show that rapid IFNγ production is dependent on glycogen breakdown (Figure 4E-F). Since glycogen is metabolized into G6P and then fed into the glycolytic machinery, we blocked the glycolytic machinery further down that pathway using the GAPDH inhibitor heptelidic acid and show diminished MAIT cell degranulation (Figure 4G). To further support the concept that glycogen supports early MAIT cell metabolic process we performed seahorse analysis on CP91149 treated MAIT cells after 3 hours of stimulation and show that early glycolysis is dependent on the breakdown of glycogen (Figure 4H-I).

**Figure 4.**
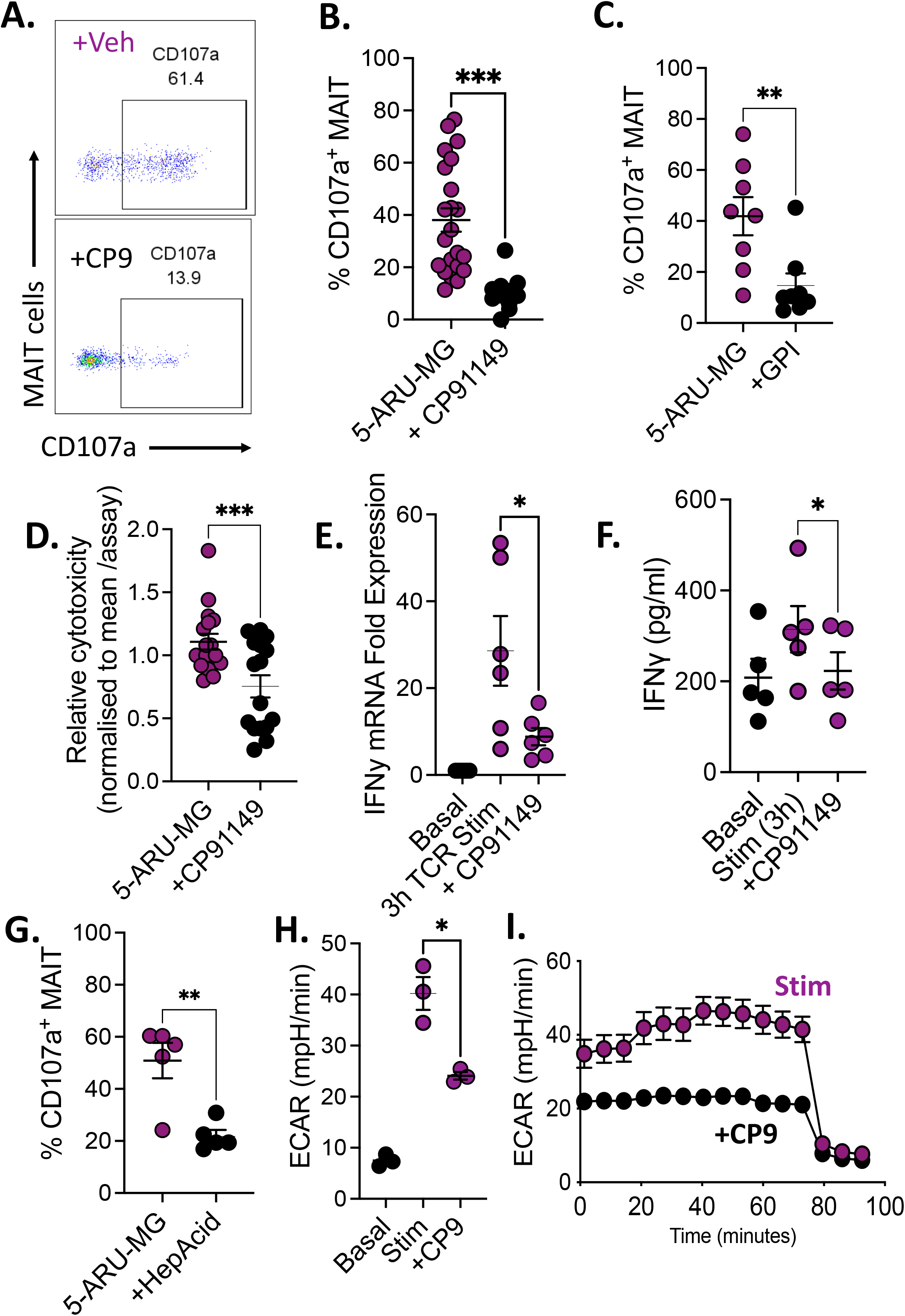
Glycogen supports MAIT cells cytotoxicity and early cytokine responses. **(A-B)** Flow cytometry dot plot and scatter plot showing CD107a expression on *ex-vivo* MAIT cells cultured with K562 cells (pulsed with 5ARU-MG) in the absence or presence of the glycogen phosphorylase inhibitor CP91149. **(C)** Scatter plot showing CD107a expression on *ex-vivo* MAIT cells cultured with K562 cells (pulsed with 5ARU-MG) in the absence or presence of the glycogen phosphorylase inhibitor GPI. **(D)** Scatterplot showing the impact of CP91149 on cytotoxic capacity of MAIT cells against K562 cells, displayed as fold change per sample. **(E-F)** Scatter plot showing IFNγ mRNA levels or secreted protein from IL-2 expanded MAIT cells stimulated with TCR beads (for 3 hours) in the absence or presence of CP91149. **(G)** Scatter plot showing CD107a expression on *ex vivo* MAIT cells in response to stimulation with K562 (pulsed with 5ARU-MG) in the absence or presence of the GAPDH inhibitor heptelidic acid. **(H-I)** Scatter plot and Seahorse trace displaying ECAR rates in TCR bead-stimulated (3 hours) IL-2 expanded MAIT cells treated with CP91149. * = p < 0.05, ** = p < 0.01, *** = p < 0.001 as measured by Wilcoxon test, paired t test or Mann-Whitney test where appropriate.

## Discussion

MAIT cells are a subset of unconventional T cells which due to their potent effector functions and abundance have been shown to play an important role in the host defence against pathogens and malignancies^1, 24^ and are now under investigation as a potential immunotherapeutic agent^15, 25^. MAIT cells have been detected in both primary cancers and metastatic sites, however they are dysfunctional, losing their anti-tumour functions^13, 17, 18, 26^. Therefore, it is critical to understand the molecular and metabolic requirements for MAIT cell anti-tumour responses.

In the current study, we confirm the robust cytotoxic potential of MAIT cells, with rapid degranulation and dose-dependent killing of K562 target cells. We also demonstrate that MAIT cell cytotoxicity of cancerous cells is dependent on MR1 and boosted in the presence of antigen, confirming work in the setting of bacterial and virally infected cells^9, 12^. Although cancer specific antigens for MAIT cells have yet to be identified^27^, the loading of cancer metabolites onto MR1 has been described^28, 29^. In addition, there is evidence emerging for microbial activation of tumour-infiltrating MAIT cells^30^.

Currently, data on the molecular regulation of MAIT cell cytotoxicity remains unclear and will be necessary as they move towards therapeutic targets. Our group and others have previously highlighted the importance of glucose metabolism for MAIT cell effector functions such as cytokine production and proliferation^20, 21, 22^. We have also reported how altered MAIT cell metabolism underpins defective functions in obesity, potentially driving pathogenic MAIT cells^20, 31, 32^. Here, we demonstrate that MAIT cell cytotoxicity (and rapid cytokine production) is independent of exogenous glucose metabolism. In our search for an alternative carbon source, we observed that MAIT cells have the molecular machinery to synthesize and metabolise glycogen. Glycogen is the main energy storage form of glucose in the body, stored as a quickly mobilised multibranched polysaccharide^33^. Recent work by Zhang and colleagues reported that murine memory CD8+ T cells but not naïve CD8+ T cells could also synthesize and metabolize glycogen^34^. We hypothesized that MAIT cells, due to their “innateness”^8^ might utilize glycogen to support their rapid functional responses. Using a series of experiments, we show that MAIT cell cytotoxicity and rapid cytokine responses (<3 hours) are dependent on the breakdown of glycogen, supporting the concept that stored glycogen fuels rapid responses in innate effector T cells like MAIT cells and memory T cells. This is further supported by work in another innate immune subset, dendritic cells, which also utilize glycogen to fuel their rapid responses^35^. Our data suggest that TCR triggering activates PYGB to break down glycogen, which then feeds glycolysis. This again is supported by the recent publication in murine memory CD8+ T cells, where the inhibition of glycolysis at the first enzyme (hexokinase) had no impact but inhibition further down the glycolytic pathway limited cellular responses^23^.

Understanding the carbon sources required to fuel MAIT cell effector functions may be of particular importance in the setting of cancer where limited glucose has been shown to impair T cell responses^36, 37^. The ability of human MAIT cells to use stored glycogen to fuel their cytotoxicity, paired with their rapid functional responses, unrestricted properties, capacity for ex-vivo expansion and relative abundance further highlights their potential as an exciting candidate for cancer immunotherapy. In conclusion, we describe for the first time a novel metabolic pathway in human MAIT cells necessary for their rapid effector responses, further supporting the rationale for their use as an immunotherapeutic.

## Materials & methods

### Study cohorts & ethical approval

Full ethical approval was obtained from both St Vincent’s University Medical Ethics Committee and Maynooth University Ethics Committee. We recruited a cohort of healthy adult donors from St Vincent’s Healthcare Group. Inclusion criteria included ability to give informed consent, 18-55 years of age and a BMI<28. Exclusion criteria included current or recent (<2 weeks) infection, current smoker, use of immunomodulatory or anti-inflammatory medications.

### Preparation of peripheral blood mononuclear cells (PBMC) and expanded MAIT cells

PBMC samples were isolated by density centrifugation over Ficoll from fresh peripheral blood samples. PBMCs were either stored at −70°C or used immediately for MAIT cell expansion using 5-ARU-MG and IL-2 as previously described^21^.

### MAIT cell degranulation assay

PBMCs were thawed and rested before addition of either metabolic inhibitors or vehicle control (i.e. 1mM 2DG, 100μM CP91149, 50μM GPI (CP316819), 5μM Heptelidic acid or DMSO / water). K562 cells with or without pre-treatment with 5-ARU-MG were then cocultured with the PBMCs at a ratio of 10:1 PBMC:K562 plus CD107a antibody (Miltenyi). After 30 minutes, protein transport inhibitor cocktail (Invitrogen) was added and the co-culture continued for a further 2 hours. MAIT cells were identified by flow cytometry with staining using specific surface monoclonal antibodies namely; CD3, CD161 and TCRVα7.2 (all Miltenyi), and degranulation assessed according to percentage of MAIT cells expressing CD107a (Miltenyi). Cell populations were acquired using a Attune NXT flow cytometer and analysed using FlowJo software (Treestar). Results were expressed as a percentage of the parent population as indicated and determined using flow minus-1 (FMO) and unstained controls.

### MAIT cell cytotoxicity assay

IL-2 expanded MAIT cells^21^ were co-cultured with Calcein AM labelled K562 cells at a ratio of 3:1 MAIT cells to target cells (and other ratios for dose curve) in the absence or presence of metabolic inhibitors or vehicle control (i.e. 1mM 2DG, 100μM CP91149, 50μM GPI (CP316819), 5μM Heptelidic acid or DMSO / water). After 2 hours of co-culture, supernatant was analysed using a BMG Clariostar multi-mode microplate reader to measure supernatant fluorescence at 485nm excitation and 525nm emission and percentage killing calculated as a proportion of maximum killing by Triton X.

### MAIT cell cytokine analysis

IFNγ mRNA and protein were measured in IL-2 expanded MAIT cells (with or without stimulation with CD3/CD28 Dynabeads (Gibco)) and their culture supernatant by qPCR or ELISA respectively. To investigate the metabolic requirements of early cytokine responses, activated MAIT cells were treated with metabolic inhibitors or vehicle control (i.e. 1mM 2DG, 100μM CP91149 or DMSO / water). mRNA was extracted from MAIT cells using Trizol according to the manufacturer’s protocol. Synthesis of cDNA was performed using qScript cDNA Synthesis kit (QuantaBio). qPCR was performed using PerfeCTa SYBR Green FastMix Reaction Mix (Green Fastmix, ROX™) (QuantaBio) and KiCqStart primer sets (Sigma). ELISAs were performed as per the manufacturer’s instructions (R&D Systems).

### MAIT cell glycogen machinery analysis

The identification of glycogen synthase (GYS-1) and phosphorylase (PYGB) in MAIT cells was based on *in silico* analysis of a published MAIT cell proteomic dataset^21^. Expression was confirmed via western blotting on expanded MAIT cells stimulated with CD3/CD28 Dynabeads for 6 hours before harvesting. Cells were lysed in NP-40 lysis buffer (50mM Tris-HCI, pH 7.4, containing 150 mM NaCl, 1% (w/v) IgePal, and complete protease inhibitor mixture (Roche)). Samples were resolved using SDS-PAGE and transferred to nitrocellulose membranes before analysis with anti-GYS-1 (Cell Signalling), PYGB (Cell Signalling) and anti-β-Actin (Sigma) antibodies. Protein bands were visualised using enhanced chemiluminescence.

### MAIT cell glycogen content and PYGB activity analysis

Glycogen content in MAIT cells (either basally or stimulated for 3 hours with CD3/CD28 Dynabeads) was measured using Biovision glycogen kit (K646) according to the manufacturer’s instructions, and by fluorescent microscopy using a previously published method^38^. Glycogen phosphorylase activity in MAIT cells (either basally or stimulated for 3 hours with CD3/CD28 Dynabeads) was measured using Sigma-Aldrich Glycogen Phosphorylase Colorimetric Assay Kit (MAK417) according to the manufacturer’s instructions.

### MAIT cell seahorse analysis

Expanded MAIT cells were treated with metabolic inhibitors or vehicle control (i.e. 1mM 2DG, 100μM CP91149, or DMSO / water) and then stimulated with CD3/CD28 Dynabeads. After 3 hours of stimulation Seahorse metabolic flux analysis was performed according to the Seahorse instruction manual.

### Statistics

Statistical analysis was completed using Graph Pad Prism 9 Software (USA). Data is expressed as mean±SEM. We determined differences between two groups using student t-test and Mann Whitney U test where appropriate. Analysis across 3 or more groups was performed using ANOVA or appropriate non-parametric test. Statistical significance was defined as p<0.05.

## Contributors Statement

FCC, NKM, EB, AB, AW and RB performed the experiments and carried out analysis and approved the final manuscript as submitted. OR and DOS recruited peripheral blood donors. AEH, DOS & FCC conceptualized and designed the study, analyzed the data, drafted the manuscript, and approved the final manuscript as submitted.

